# Importance of agriculture for Crop Wild Relatives conservation in Switzerland

**DOI:** 10.1101/2022.10.05.511054

**Authors:** Blaise Petitpierre, Julie Boserup, Adrian Möhl, Sibyl Römetsch, Sylvain Aubry

## Abstract

Crop Wild Relatives are a subset of the global plant diversity that is often neglected, as not the primary focus for conservationists or plant breeders. However, a relatively large portion of the wild flora, up to 60% in Switzerland for example, do share genetic relationships with cultivated species and therefore can be considered as Crop Wild Relatives. Their conservation appears simultaneously a challenge to conservation programmes but also a considerable levy to mobilize other sectors, like agriculture, to contribute to the conservation of biodiversity at large. Here, we provide a comprehensive checklist of Swiss Crop Wild Relatives representing 2,226 taxa, of which 285 prioritised taxa, referred to as “Crop wild relatives Of Concern”, were designated. Following a taxa-specific ecogeographic analysis, we analysed the extent to which CWR of concern are already contained in existing protected areas as well as their distribution in the agricultural area. Prioritised Crop Wild Relatives species richness was compared to modelled species richness to identify potential conservation gaps. About a fifth of CWR of concern is not significantly better protected than a random species by existing protected areas. However, 28.8 % and 15.5 % of these taxa are more frequently distributed in agricultural and summer grazing areas respectively than random expectations. A clear deficit of species richness for these Crop Wild Relatives of concern was inferred on low lands, possibly related to a lower sampling effort. We further identified a network of 39 sites that contains all taxa of Swiss CWR of concern and that could be used as a primary conservation infrastructure. More generally, our results could be generalized to other countries and support better consideration of CWR in agriculture areas, an important “reservoir” for expanding specific measures of conservation that are crucial to meet the future global goals of diversity conservation frameworks.

## Introduction

Crop diversity, across and within species, is a major driver of agricultural resilience. However, it is estimated that seventy-five per cent of crop diversity was lost globally during the 20^th^ century (FAO 2010). The loss of allelic diversity in crops is partly inherent to the breeding process but also due to a wide range of other socio-economic factors that gradually led to the narrowing genetic basis used for breeding (Hajjar & Hodgkin 2007; Khoury et al. 2022). Indeed, most crops originated from the domestication of wild ancestors (Engels & Thormann 2020). Therefore, attempts to identify and save Crop Wild Relatives (CWR, Maxted et al. 2006) appear as a priority to ensure a viable future for the next generations of farmers and breeders and eventually improve food security. More generally, CWRs often represent a very large portion of the wild flora (Maxted et al. 2006) and a better focus on their conservation may actually raise unexplored levies, that in turn could be essential for biodiversity conservation at large.

Many breeding programs increase their genetic basis by integrating CWR: for example, the resistance to late blight (*Phytophthora infestans*) from the wild potato *Solanum demissum* or the stem rust resistance (*Puccinia graminis*) from wheat’s CWR *Aegilops tauschii* (Hajjar & Hodgkin 2007; Dempewolf et al. 2017). Since the CWR concept has been described in the seventies, growing international momentum for their conservation has emerged, side-by-side with the efforts aimed at global biodiversity conservation (Harlan & Wet 1971; Maxted et al. 2006). Recently, global (Maxted et al. 2012; Castañeda-Álvarez et al. 2016; Vincent et al. 2019) as well as national (Rubio Teso et al. 2021) inventories of CWR revealed an urgent need for measures to preserve CWR diversity. Many countries performed their own CWR inventory, for example, Portugal (Brehm et al. 2008), Norway (Phillips et al. 2016), the Czech Republic (Taylor et al. 2017), the UK (Jarvis et al. 2015), Netherland (Treuren et al. 2017), Turkey (Tas et al. 2019), USA (Khoury et al. 2013), mostly pursuing similar aims but slightly divergent methodologies. The procedure typically comprises several successive steps: inventory, prioritisation and identification of potential “hot spots” for their conservation, often referred to as gap analysis (Maxted et al. 2007). In addition, metrics representative of the relative importance of crops for agriculture could be scored, as it has been performed globally, to inform food security policy (Castañeda-Álvarez et al. 2016).

In Switzerland, a national action plan to conserve crops *in situ* and *ex situ* has been implemented since 1999. It is not primarily focusing on CWR and has only partially succeeded in counteracting the decrease in agrobiodiversity (Guntern et al. 2013). Swiss agriculture covers about a third of the country’s surface and therefore represents major pressure on ecosystems (FOAG & FOEN 2013; Guntern et al. 2013). The most recent data showed that more than half of the habitats and 36% of all species are threatened or near-threatened (Guntern et al. 2013). Interestingly, Swiss farmers are entitled to public subsidies, in the form of direct payment following a “cross-compliance” scheme. These subsidies are conditioned to a set of practices and provide proof of ecological performance (Jarrett & Moser 2013). Briefly, this entails limited fertilization and pesticide use, crop rotation, animal welfare measures and 7% of the land allocated as ecological compensation area (ECA). Based on the overwhelming influence of cultivated areas on wild ecosystems, we wondered how prevalent the agricultural areas, their various mode of management and their associated public subsidies could underpin CWR and more generally plants conservation. To answer this question, we first built a comprehensive checklist of CWR in Switzerland using a floristic approach (Maxted et al. 2006). We then derived a list of “CWR of Concern” (CoC) taxa. The CoC concept allows us to effectively merge conservation and utilization concerns without taking the risk of confusion with the generally used “priority/prioritisation” for threatened species. The list of CoC contains therefore either CWR that need conservation measures and CWR that are very closely related to cultivated crops. We then performed an extensive ecogeographical analysis of CoC using observed and modelled species distributions. Subsequently, we evaluated the extent to which CoC are already protected under various “conventional” measures (parks and other protected areas). Finally, we measured the actual overlap between CoC distribution, the agricultural land, their dedicated ECA and identified a minimal network of sites that include all CoC across Switzerland.

Combining data on the state of CWR conservation and their overlap with agricultural areas in Switzerland, we draw some conclusions on ways to improve conservation policies. We believe our approach could be generalised to other countries and can improve consideration of the link between land management and public action in sustaining CWR conservation and biodiversity more generally.

## Methods

### List of crops with relevant use in Swiss agriculture

A list of 129 crops for food and feed that are grown in Switzerland was built (Appendix S1). It contains major and minor crops as defined by Khoury et al. (Khoury et al. 2013). This comprises all species from Annex 1 of the International Treaty for Plant Genetic Resources for Food and Agriculture and all crops contained in the Swiss national databank for cultivated plants (FOAG 2022) and trade data. In addition, a list of the most common forage species was obtained using the latest Swiss forage crop recommendations (Suter et al. 2019) as well as a list of modern medicinal plants from a published ethnobotanical survey (Cero et al. 2014). Species cultivated as aromatics were collated with help from local experts.

### Swiss CWR checklist and prioritization

A CWR checklist was created for Switzerland primarily based on information from the Crop Wild Relative Catalogue for Europe and the Mediterranean (Kell et al. 2008, updated version by Maxted & Kell, personal communication). Using a floristic approach, taxa information was updated and harmonized with the latest version of the checklist of the Swiss Flora published (Infoflora 2017).

We then linked every possible taxon out of the 2,226 CWR from the checklist with a given utilization, based on our initial crop list (Appendix S1), namely primary or secondary food, forage, medicinal, aromatic, industrial, restoration, forestry and ornamental (Table 2 and Appendix S2). We considered any species contained in a genus having at least one reported use was considered as a CWR (or a wild-used species, no distinction has been made). Compiling data from published checklists from other European countries and the US (Brehm et al. 2008; Khoury et al. 2013; Fielder et al. 2015; Phillips et al. 2016; Taylor et al. 2017; García et al. 2017), we analysed the extent of the relationship between crops and their respective CWR, defined by either the Gene Pool (GP) and the Taxon Group (TG) concepts (Harlan & Wet 1971; Maxted et al. 2006).

**Table 1.** Checklist of CWR in Switzerland. A checklist of CWR has been realised merging data from the European CWR list and the *Flora helvetica* latest version (2017). Only CWR related to cultivated crops (not considering ornamentals) have been considered further for prioritization.

|  | **No of CWR taxa** |
| --- | --- |
| Swiss CWR according to CWRIS (Kell et al., 2005) | 4'464 |
| Swiss CWR checklist after correction using *Flora helvetica* (Infoflora, 2017) | 3'006<br>(60% Flora) |
| **Swiss CWR checklist** (without neophytes and invasive species and without taxa related to ornamentals) | **2'226** |
| CWR from PGRFA | 2'045 |
| **CWR of Concern (incl. expert's opinion, see Methods)** | **285** |
| CWR of Concern in the red list 2016 | 90 |
| CWR of Concern in the list of the National Priority Species (FOEN) | 92 |

**Table 2.** List of CWR taxa according to their reported use. This is based on the 2,226 taxa from the CWR checklist (Appendix S2). For each category, the number of taxa is indicated, as well as their degree of relationship (Documented Gene Pool or Taxon Group) and their status of conservation on the red list (IUCN, 2016). Note that some taxa can belong simultaneously to several categories.

|  | Food | Forage | Medicinal | Aromatic | Industrial<br>Restoration<br>Forestry |
| --- | --- | --- | --- | --- | --- |
| **No of CWR taxa** | 334 | 211 | 1438 | 28 | 114 |
| **Documented**<br>**relationship** | 140 | 119 | 497 | 3 | 34 |
| **Conservation status**<br>**(CR, EN, VU, DD)** | 62 | 30 | 300 | 2 | 27 |

We prioritized a shorter list of CoC, for which conservation measures should be ensured to maintain genetic diversity among populations. Four criteria have been selected and scoring applied on a scale from 1 (low priority) to 5 (high priority): 1. if a known relationship of the CWR to a currently cultivated crop in Switzerland was described, a high priority score was given; 2. if a CWR had a close relationship to a crop, a higher score was attributed (GP1/TG1>TG/GP2>TG3>TG4); 3. conservation status was assigned based on the IUCN red list classification, with 5 points to taxa classified as critically endangered (CR), 4 to endangered (EN), 3 to vulnerable (VU), 2 to near threatened (NT) and 1 to least concerned (LC); 4. Finally, the origin of the taxa was taken into consideration, with a maximum priority score (5) for indigenous and archeophytes, a score of 2 for European neophytes, and a score of 1 for neophytes. Finally, the list of CoC was submitted to expert advice that took into consideration interests for breeding and use in Swiss agriculture. Results from the first expert consultation performed by Häner et al (Häner et al. 2009) were also compiled and integrated.

### Species distribution, protected area and agricultural surface

To identify species richness, observations of CoC recorded between 01/01/2002 and 31/12/2019 were extracted from the database of the Swiss national data centre for vascular plants (Infoflora, 2020). Cultivated or sub-spontaneous occurrences were removed, so as occurrences with an uncertainty > 250 m. To avoid duplication, species observations were disaggregated keeping a minimal distance of 100 m between occurrences. In total, 567’319 observations were used in the analysis.

To summarize a comprehensive set of protected areas over the country we combined the geographical layers of Federal inventories (FOEN 2018), the natural reserves managed by Pro Natura, forest reserve (FOEN 2018) and the Swiss National Park were bulked together. The agricultural surface has been determined using data from (Szerencsits et al. 2018), with a distinction between the actual agriculture surface and the surfaces dedicated to summer grazing being retained.

To assess if the distribution of each CoC was significantly overlapping the protected areas, we generated 1,000 random distributions. These random distributions consist of 1’991 points (corresponding to the average number of occurrences among the CoC), sampled following the sampling bias found in the Infoflora database (Fig. 1). For each randomization, we measured the proportion of the random distributions covered by the protected area, allowing us to test if the taxa were significantly more protected by the existing protection area than expected by chance.

**Figure 1.**
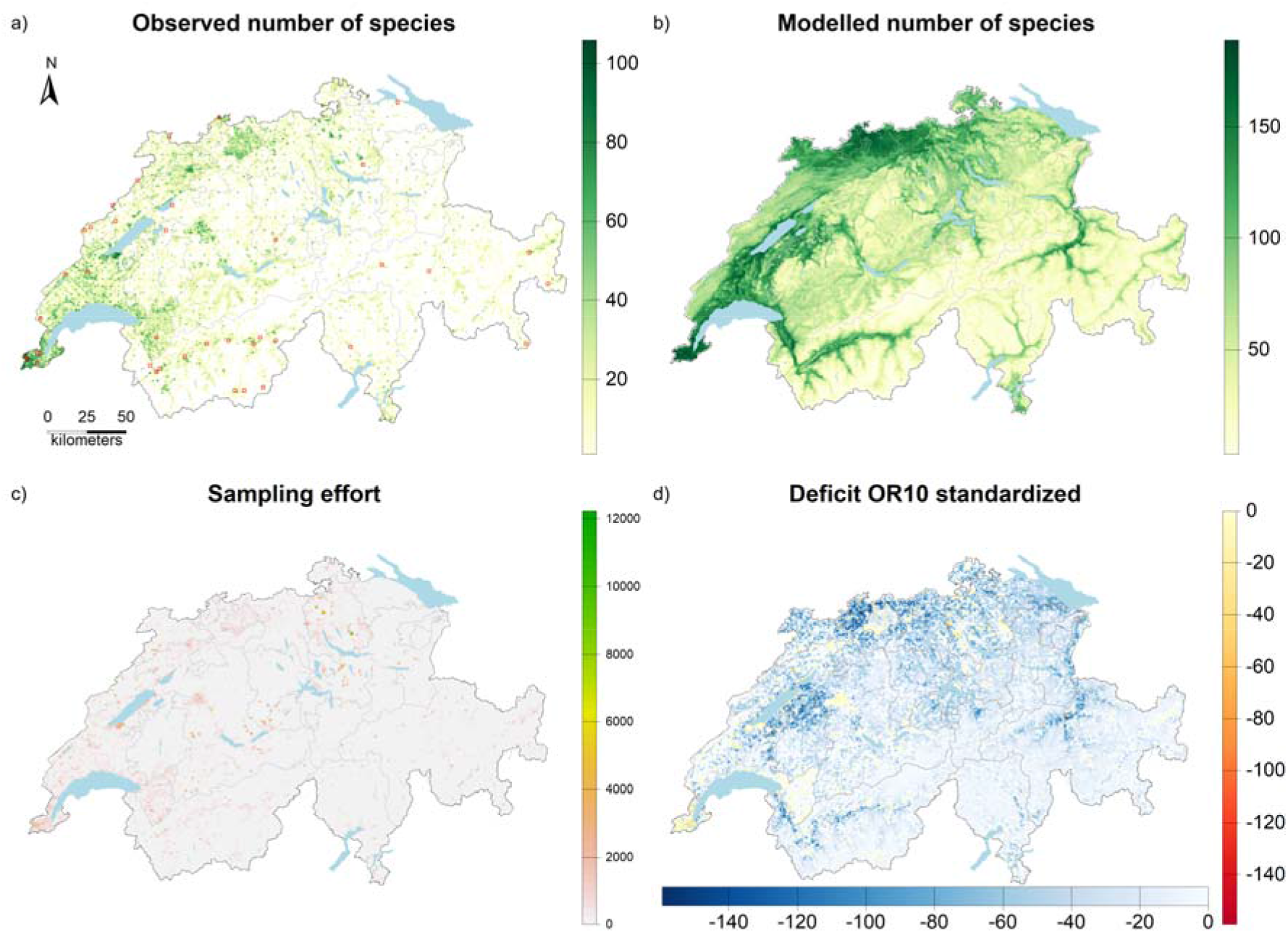
**a)**. CWR of concern (CoC) were observed in Switzerland. Red sites represent the minimum number of sites to cover all CWR species in the country. b) Modelled distribution of CoC in Switzerland obtained by the stacking of the potential distribution of each priority CWR in Switzerland. c) Sampling effort represented by the number of observations for all plant species in the Infoflora database for the period 2001-2019. d) Deficit area between observed and modelled distribution of CoC in areas with a high (yellow to red) or lower sampling effort (light to dark blue).

### Species distribution modelling

Species distribution models (SDMs) relate species occurrences to environmental factors. Once this relation is statistically quantified, it is then possible to derive predictions of species potential distributions if the predictors are spatially explicit (Guisan & Thuiller 2005; Elith & Leathwick 2009). SDMs are particularly useful for conservation practices (Guisan et al. 2013). In this study, we built potential distribution maps derived from SDMs for every taxon with enough observations (n = 10). For each species, predictors were selected from an initial set of 33 variables including information about the topography, climate, soil and remote sensing (Appendix S3). A preliminary variable selection was processed for each taxon to reduce the number of predictors and avoid model overfitting. After this initial step, the number of variables varied between two and nine, depending on the species (Appendix S4). These predictors were related to species occurrences by combining three modelling algorithms (general additive models, Maxent and gradient boosting model) into an ensemble modelling approach (Thuiller et al. 2004; Araujo & New 2007), or an ensemble of small models (ESM, Breiner et al. 2015) depending on the number of observations (Appendix S4). Models were evaluated with 4-fold cross-validation with an index combining 4 commonly used indices of accuracy (AUC, TSS_max_, Sensitivity and continuous Boyce index, Appendix S4). This index is analogous to a correlation varying between −1 (total counter predictions) and 1 (perfect predictions), 0 meaning random predictions.

### Patterns of species richness

We estimated the difference between the modelled and the observed species richness to map the deficit between the observed and the modelled number of CoC species. Because the stacking of SDM maps is known to be sensitive to the threshold used to binarize continuous suitability maps (Benito et al. 2013; Calabrese et al. 2014; Schmitt et al. 2017), we applied five different thresholding criteria to reclassify the individual species suitability maps into potential presences and absences (Appendix S4). As modelled species richness obtained by stacking SDM maps tends to be overestimated (Guisan & Rahbek 2011; Calabrese et al. 2014), we applied a quantile normalization between the map of the observed number of species and the modelled number of species. Quantile normalization was initially developed for the analysis of high-throughput data in molecular biology (Amaratunga & Cabrera 2001; Bolstad et al. 2003). In our case, it allows standardizing all the distributions of richness (observed and modelled) with the same minimal and maximal values, while keeping their statistical properties (Hicks & Irizarry 2015). Finally, we included the sampling effort to interpret the deficit between modelled and observed distributions. We gathered all observations for all the plant taxa recorded in the database of Infoflora between 2001 and 2019 and categorized areas with a high (≥ 500 observations per km^2^) and low sampling effort (respectively ≥ 500 and < 500 observations per km^2^).

### Complementary analysis to delimit a minimal conservation network

A complementary analysis was carried out to obtain a spatial network that most efficiently covers CoC species. We selected the site with the highest number of taxa, excluded these taxa from the analysis and iteratively repeated this process until all taxa were covered (Rebelos 2014). This analysis was applied to the observations of the 285 CoC species distributed on a 4 km^2^ grid. All the data analysis was run with a custom R script available on demand (version 4.0.3; R Core Team 2020).

## Results

### Swiss CWR checklist and prioritization

A total of 3,006 taxa were identified as CWR, representing about 60% of the described Swiss flora (Table 1). Among those, taxa classified as invasive neophytes species as well as taxa related to ornamentals were removed, leaving 2,226 CWR, of which 2,045 related to any agricultural use (namely food, feed, medicinal, aromatic, restoration). For prioritization purposes, ornamentals were removed as they represent a very large portion of the flora. Noteworthy, while the conservation status of various taxa has been extensively documented (Red list, priority species), only a relatively small proportion of CWR relationships have been reported. For example, information about the genetic relationship to their crop (gene pool) could only be documented from the literature for 140 out of 340 CWR taxa of food crops (Appendix S2).

Based on four criteria: the relationship to a species used in the Swiss agroecosystem, the genetic distance to a specific crop, its conservation status and its origin, 285 CWR were considered as “CWR of concern”. Importantly, we also considered 18 CWR of crops not grown in Switzerland, like *Setaria*, and *Hedysarum*). While Switzerland does not hold formal responsibility for such taxa, safeguarding some of the genepools that may be useful for other agrosystems seems reasonable in view of global climate change. To validate this list, we compiled expert data available (Häner et al., 2009) and new data to evaluate the relevance of this list to breeding and conservation sectors. This allowed, for example, to integrate into the CoC list, species like *Artemisia annua L*. or *Rhodiola rosea L*., where local research for medicinal application and breeding has been undergone (Simonnet et al. 2008; Vouillamoz et al. 2012). Among the 257 taxa (out of 285 CoC) with a national red list conservation status, 148 are least-concerned (LC; 51.9%), 21 near-threatened (NT; 7.4%), 49 vulnerable (VU; 17.2%), 24 endangered (EN; 8.4%) and 15 critically endangered (CR; 5.3%) taxa. Among the CoC, 92 taxa (32.3 %) taxa belong to the list of the National Priority Species requiring conservation measures (FOEN 2019).

### Distribution, richness and deficit areas of priority CWR in situ

The areas with the highest CWR taxa richness were found in the northwest region of the country at relatively lower altitudes (Figure 1a). The observed richness is correlated with the sampling effort (Spearman correlation between the number of observations and species richness at a 1km^2^ resolution: rs = 0.799, n = 39’961, P < 0.001; Figure 1c).

SDMs were generated for 265 of the 285 priority CWR (Appendix S5). For 20 taxa, the reduced number of observations available from the database (< 10) could not generate reliable modelling. These 20 taxa are composed of 10 known rare species covered by the national Red List belonging to the National Priority Species list, 7 subspecies requiring expert knowledge to reach this determination level and 3 taxa with very few documented observations in Switzerland. The consensus evaluation index varies between 0.4 and 0.968 (with an average of 0.774), supporting that the modelled distributions are accurate. For each CoC taxa, like for example *Allium lineare* (Figure 2b), maps representing observations, habitat suitability and potential distributions were generated (Appendix S5). For *Allium lineare*, clear potential distribution was flagged in Wallis and Graubünden (Figure 2c & d). Only 10 species were modelled with an accuracy below 0.6 (Appendix S6). These 10 taxa were removed from the analysis of the deficit, in addition to the 20 taxa with insufficient observations. Therefore, the comparison between observed and modelled distributions of the species richness was done with 255 species accurately modelled.

**Figure 2.**
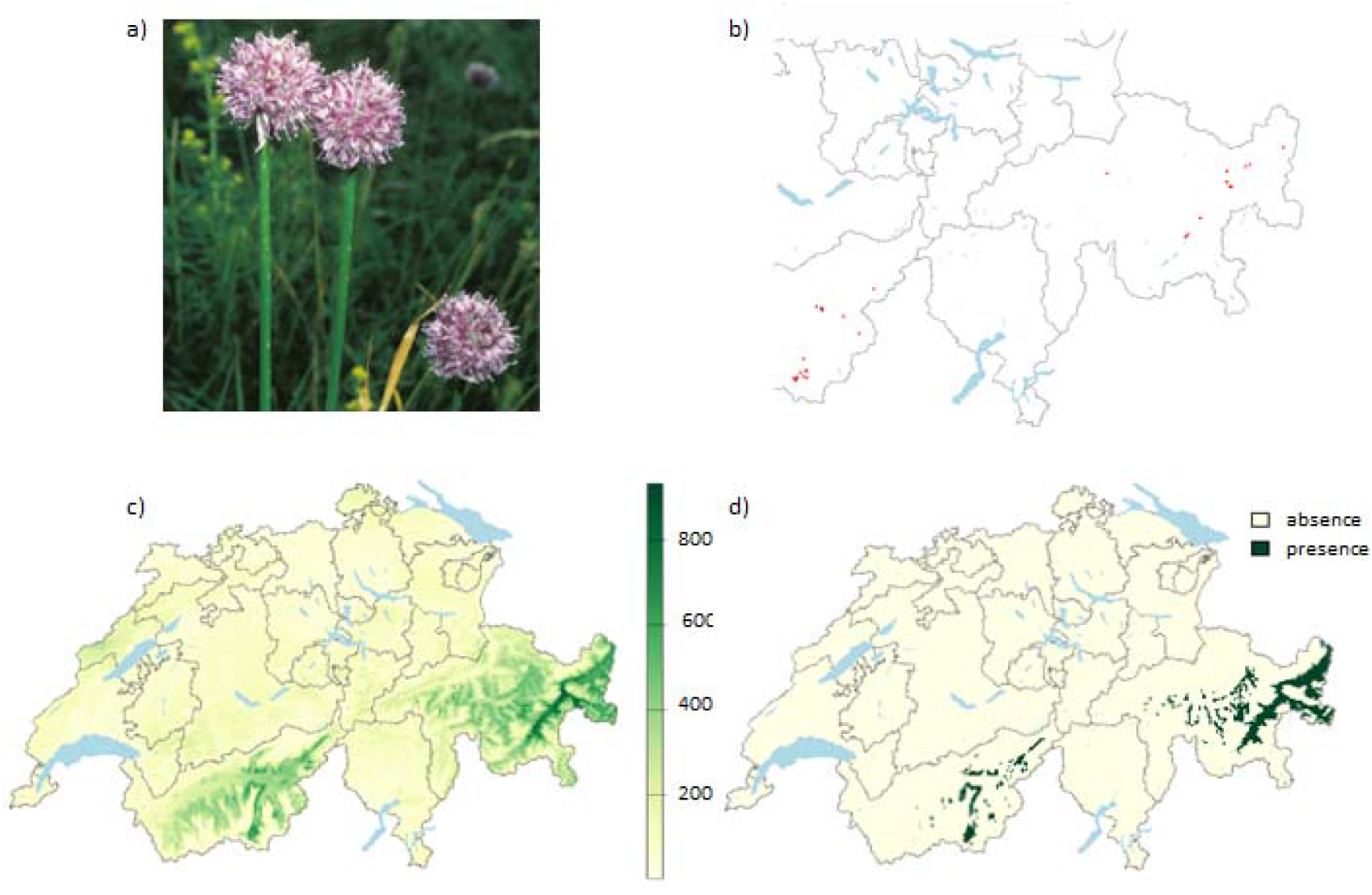
*Allium lineare*, a CoC from onion (a) and its known distribution in Switzerland (b; each red dot is a single observation). Species distribution models produce continuous habitat suitability maps (c), which are binarised into a potential distribution map (d). Here, the omission ratio of 10 (OR10) was used but 4 others thresholding criteria were used (Appendix S6)

Not surprisingly, the modelled and observed distribution of CoC are correlated (Spearman correlation between observed and modelled species richness at a 1km^2^ resolution: average rs across the 5 thresholding methods = 0.491 ± 0.016, n = 39’961, P < 0.001; Figure 1a and b). The modelled richness correlates with the sampling effort much less than the observed species richness (Spearman correlation between the number of observations and the modelled species richness at a 1km^2^ resolution: average rs across the 5 thresholding methods 0.412 ± 0.019, n = 39’961, P < 0.001, Figure 1).

The comparison between the observed and potential species richness shows an important deficit at the lowland elevation, with some obvious gaps in regions like the Swiss Plateau, Wallis, Ticino and Graubünden. However, it appears that most of this deficit appears in areas with a low sampling effort. In areas with higher sampling effort, there is less deficit (Figure 1d).

### Distribution of CWR of concern in protected and agricultural areas

On average, CoC have 33 ± 21.4 % of their distribution located within protected areas (Table 3). This is significantly more than the distributions of the null model (13.9 ± 0.1 %; p-val. of a two-sample t-test < 0.001; Table 3). However, 64 species (22.5 % of the CoC) are not significantly more protected than a random species (Appendix S7). Taking advantage of our data, we considered further the probability for CoC, as distant relatives of crop plants, to share some habitats with cultivated plants in the agricultural or summer grazing areas. The eco-geographical analysis reveals that on average 20.1 ± 15.3 % of the CoC are located within the agricultural area (Table 3). This is not significantly more than the distributions of the null model (26 ± 1 %; p-val of a two-sample t-test = 1; Table 3) but it is noticeable that 82 species (28.8 % of the CoC) are significantly more distributed in the agricultural area than expected by chance (Appendix S7). Finally, CoC are not significantly more distributed in summer grazing areas. On average, 4.8 ± 7.8 % of their distribution is located within summer grazing areas, whereas 9.2 ± 0.6 % of the random distributions fall within summer grazing areas (p-value of a two-sample t-test = 1; Table 3). Nevertheless, 43 species (15.1 %) are significantly more present in summer grazing areas than expected by chance (Appendix S7).

**Table 3.**
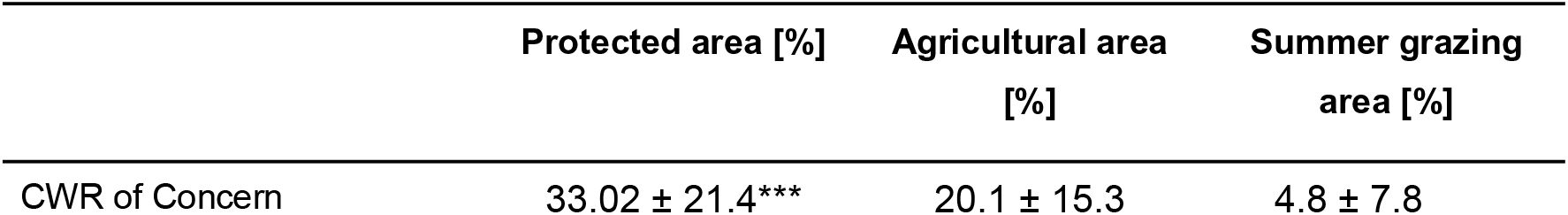
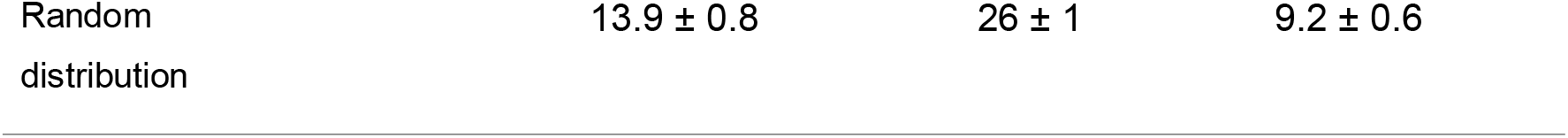
Average distribution of CWR of Concern in protected, agricultural and summer grazing areas. For comparison, we also provide the average distribution of the 1’000 random distributions of the null model. *** significantly more than randomly distributed (p-val < 0.001).

### Minimal conservation network of CoC for an adapted in situ conservation

The minimal spatial network to cover at least one population of all the 285 CWR taxa consists of 39 2-by-2 kilometres squares, mostly located in South-western Switzerland (Figure 1 a). In these 39 sites, the proportion of protected area ranges from 0% to 51%, with an average of 8% (Appendix S8). The proportion of agricultural and summer grazing areas dedicated to summer grazing in these “hotspots” ranges from 0.8 to 86%, with an average of 35.2 % (Appendix S8). Interestingly, the proportion of protected area within the hotspots is not correlated with the proportion of agricultural area (Pearson’s correlation P = 0.029; p-val = 0.862), neither with the summer grazing (Pearson’s correlation P = −0.071; p-val = 0.668).

## Discussion

### Adapting conservation priorities in a changing environment: the Swiss CWR inventory

Following a global effort to improve the conservation effort of CWR globally (Vincent et al. 2013), we took advantage of the recently updated checklist of the Swiss flora (Infoflora 2017) to generate a comprehensive country-wide CWR inventory. With an overwhelming 60% of its entire flora being considered as CWR, including a significant number of plants relative to medicinal plants (1,438, Table 2), this checklist had to be prioritized. This process identifies CoC and allows a dedicated set of measures depending on their respective conservation status: while the most vulnerable taxa are or will be included in current conservation plans, other less threatened taxa may benefit from some monitoring of their populations. Combining four sets of criteria (relationship to cultivated species, degree of relationship, red-list status and origin of the taxa) and validated by experts, we short-listed 285 CoC taxa that will be targeted by various dedicated measures, depending on their conservation status. Logically, our priority list partially overlaps with the recently published European priority list (Rubio Teso et al. 2021).

While the IUCN red list and the list of the National Priority Species are key elements to identify taxa that are immediately threatened and will be central to identifying CoC that are in need of active conservation measures. But the list of CoC also contains taxa that are not threatened, while important from a breeding perspective. For example, *Daucus carota* or the various *Festuca* (*rubra, pratensis, ovina*…), are not particularly threatened according to our ecogeographical analysis (Appendix S6). However, these taxa remain an important target for CWR conservation to maintain genetic diversity within the genera of relevant crops (Khoury et al. 2022). In any case, the modularity of our approach may allow reshuffling the priority criteria and easily generating a new priority list that may be better suited for other stakeholders. This work provides the first step towards considering a portion of the biodiversity, namely the relatives to cultivated plants and wild-used plants as a valuable target for conservation policies.

### Filling the gaps in existing conservation measures to include CWR of Concern

We then used ecogeographic tools to conduct a nationwide assessment of the conservation gaps for each of the CoC. We first assessed the extent of protection of CoC in existing protected areas. Our analysis shows that although the majority of the priority taxa is well covered by existing protected areas, 22.5 % of CoC are not significantly better protected than a randomly distributed species. It is obvious that for some of these species, prioritization was mostly due to their close relationship to cultivated crops rather than their conservation status (e.g. *Capsella bursa-pastoris*, *Lactuca serriola*, *Lolium multiflorum*). However, this list also reveals taxa that are threatened but currently not well covered by existing protection areas (e.g. *Allium rotundum*, *Alopecurus geniculatus*, *Chenopodium vulvaria*, *Fragaria moschata*, *Lactuca saligna*, *Taraxacum pacheri*). These species are to be found in habitats that are usually not covered by habitat inventories. For example, most of the dry meadows in Switzerland are in the “dry meadows and pasture” federal inventory where they profit from adequate protection despite occurring usually in environments that are outside protected areas. This conservation gap might be due to the protected areas not necessarily targeting the CWR specifically or to an overall limited distribution and efficiency of the existing protected areas (Guntern et al. 2013). Biodiversity loss is severe in Switzerland, which is far from reaching Aichi’s targets (initially aimed for 2020, FOEN 2017). Currently, protected areas cover only 12.5 % of the country (FOEN 2017). More efforts have to be performed in the protection of natural habitats in general, including the CoC species which are already covered by the current network of protected areas. Globally, similar trends have been observed for red list plants “used for human food”, with only 47% not covered by protected areas (FAO 2010). The protection gap observed for CoC might therefore benefit from more dedicated actions, like the identification of hotspots relevant for *in situ* conservation. Several successful examples of CWR-specific protected areas have been documented, while issues related to the required standards and conflicts with local land management policies were reported (Iorondo et al., 2012, Fielder et al., 2015). Based on the rationale that CWR might share, to a certain extent, similar ecogeographic zones with the crop they are related to, we concentrate next on the potential for the agricultural areas to be better considered in the CWR conservation strategy.

### Agriculture land management as levies for CWR conservation

A large portion of the CoC populations were found in relatively lower altitudes, as well as their respective deficit regions. The deficit in floristic quality could be confirmed in the lower and intensively cultivated areas compared to higher lands, as observed before (Meier et al. 2021). However, our analysis also shows that areas with a higher sampling effort have much less deficit of CoC species richness, suggesting that this apparent deficit might be also due to a lack of observation data.

Comparing observed and modelled stacked species distributions is sensitive because stacking potential distribution is known to systematically overpredict species’ richness (Benito et al. 2013; Calabrese et al. 2014; Schmitt et al. 2017). Likewise, this would artificially increase the modelled deficit. Our study shows that the amount of modelled species richness is highly dependent on the choice of the threshold used to binarise the species’ continuous suitability map (Appendix S3 and S4). However, the distribution pattern of the modelled species’ richness remains stable across the thresholding strategies (Appendix S3 and 4). This suggests that a standardisation such as quantile normalization to rescale the modelled species’ richness with the observed one, might provide a simple and conservative approach to map areas with an important difference between observed and modelled richness due to methodological bias.

Based on these observations, and again on the assumption that species that are related to cultivated species might share their ecological niche, we wanted to evaluate the extent to which the CoC populations were localized on the agricultural land. To our knowledge, there have been few (if any) attempts to try to evaluate the overlap between cultivated lands and CWR. Our analysis shows that 43.9 % of the CoC are more frequently distributed in agricultural or summer grazing areas. This important fraction supports that CWR could be an element to be integrated into the complex set of measures dedicated to the ecological compensation areas to promote farmland biodiversity (Aviron et al. 2009). The current analysis identified an interesting group of taxa, which shows on one hand bad coverage by protected areas, and on the other hand a significant part of their distribution in agricultural or summer grazing areas. For these 30 taxa (10.5 % of the CoC), agricultural measures and policies may help to better conserve these species. For example, *Valerianella dentata* is a characteristic cornfield plant found on lighter, more calcareous arable land, particularly overlying chalk (Appendix S5). Even though this species has a wide global distribution, its populations have diminished considerably and it is considered a vulnerable (VU) species on the national Red List (Bornand et al 2016). This decline has been a result of the intensive use of herbicide and the application of nitrogenous fertilizers to highly competitive modern crop varieties. It has been shown that the species can be successfully aided by adequate management of wheat fields and it is, therefore, a perfect example of an endangered species and close relative to a widely used crop.

In Switzerland, since 2018, a new *ad hoc* plan promotes *in situ* conservation of some forage crop populations. These measures target an overall surface of 2,750 ha under a dedicated cross-compliance scheme and target specifically 24 CoC. Interestingly, when considering forage plants, some conflicting aims could be identified, namely between the short-listing performed by the botanists and the farmer’s priorities. For example, *Poa trivialis* listed here as a CoC, is also considered a common weed of grazing surfaces by many farmers. This is a practical case that shows the strong need to guarantee the inclusion of all relevant stakeholders in the process of designing CWR conservation policy.

To better target potential conservation plans locally, in the last step of our analysis, we identified a network of 39 sites all over the country that allows a comprehensive coverage of all 285 CoC (Fig. 1a). Interestingly, the majority of these sites are localized in the hotter and drier climate of Switzerland optimal for agriculture (Holzkämper et al. 2015), suggesting a particularly promising area for implementation of further measures. Again here, about one-third of this conservation network is in agricultural or summer grazing areas, supporting that these surfaces are critical for an efficient conservation strategy of CWR. Because our distribution dataset mainly relies on opportunistic observations without any sampling design, the distribution of this network might be sensitive to the distribution of the sampling effort. If novel areas get better sampled, this can possibly affect the distribution of rare CoC and therefore modify the distribution of this complementary network. Such a network dedicated to *in situ* conservation of CoC could integrate information on the modelled species distribution (Guisan et al. 2013; Tulloch et al. 2016) to be less exposed to the influence of sampling effort. Another advantage of SDMs is the possible inclusion of climate or land use scenarios to project potential distributions in the future so that conservation networks could anticipate future distributional changes (Faleiro et al. 2013; Mateo et al. 2019). The current analysis can be used as a first step to synthesise current knowledge about CoC. Combining prospective field campaigns and potential distribution analyses integrating global change scenarios would inform how to complete current national monitoring such as the Swiss Biodiversity Monitoring (FOEN 2014) or the Agricultural Species and Habitats Monitoring Programme (Riedel et al. 2018) to develop efficient monitoring of the CoC.

### Raising synergies between conservation and agriculture

The objective of the current study was to set the ground for a comprehensive and sustainable strategy for CWR conservation in Switzerland. Conservation of CWR remains a “grey zone” as much for conservationists as for farmers or policymakers. If we are to meet the United Nations Convention on Biological Diversity global strategy for plant conservation, which states in its objective n°9, that “70 per cent of genetic diversity crops, including their wild relatives and other socio-economically valuable plants species {should be} conserved” (Convention for Biological Diversity 2011), a synergy between sectors appears urgent. The significant enrichment of CoC on the agricultural surface may be a specificity of the Swiss landscape, and the extent to which this can be extrapolated to other agroecosystems remains to be determined. However, the interaction between CWR and agriculture appears largely unexplored. Addressing the rapid loss of biodiversity in the near future, that in turn may directly impact our agroecosystem resilience, will require a cross-sectoral approach (Frison et al. 2011). We believe we provide here a compelling example of how the CWR conservation is a particularly good first stepping stone for enhancing synergies between agriculture and conservation.

## Acknowledgements

This study has been funded by the Swiss Federal Office for Agriculture (grant n° 05-NAP-P57). We would like to thank Raphael Häner, Jérôme Frey for helpful discussion; Andreas Gygax and Stefan Eggenberg for facilitating access to the floristic data.

## Supporting Information

**Appendix S1. Checklist of crops used in Switzerland**. Compiled list of crops for food and feed, aromatic, medicinal and ornamentals. Note that ornamental crops and their CWR were not used in the final CWR prioritization.

**Appendix S2. Checklist of Crop Wild Relatives (CWR) in Switzerland and prioritized CWR list**. A detailed list and nomenclature of the 2,226 CWR identified in the Swiss flora and the 285 CWR of Concern that have been short-listed in the current study.

**Appendix S3. List of the available environmental variables available for SDMS**. Each variable was assigned to a category: temperature (T), seasonality (S), extreme temperature (Tex), precipitation (P), aridity (A), topography (Topo), NDVI, forest height (Forest) and soil pH (pH). Each variable has a priority rank within its category for the preselection procedure. This procedure used a permutation test for which a minimum p-value (p-val th.) was required to keep the variable in the procedure.

**Appendix S4. Detailed method for SDMs.**

**Appendix S5. Actual and potential distributions for each taxa. Data available under the link:** <u>10.6084/m9.figshare.21019963</u>

**Appendix S6. Results of the SDMs for CoC**. For each species, the number of observations used to calibrate the models (Nobs), the modelling type (ESM for an ensemble of small distribution models or EM for ensemble models) and different model evaluation indices are given. In orange, models with a “not-so-good” evaluation.

**Appendix S7. Results of the eco-geographical analysis for each CoC**. For each species we provide the number of observations used to describe the species’ distribution, the IUCN red list status, the priority in the list of the National Priority Species (priority in NPS, see legend in Appendix S2), the proportion of the distribution located in protected area (protected [%]), unprotected surfaces in agricultural area (unprotected in AA [%]) and in unprotected surfaces in summer-grazing area (unprotected in SGA [%]). The distribution of each priority CWR has been compared with 1000 random distributions to estimate if the distribution was more frequent in protected area (rand. test reserve [p-val.]), in unprotected surfaces in agricultural area (rand. test AA [p-val.]) and in unprotected surfaces in summer-grazing area (rand.test SGA [p-val.])

**Appendix S8. List of 39 “CWR hotspots” that cover comprehensively the 285 CWR of Concern distribution.**

