## Appendix S3, S4, S9_Methods for "Importance of agriculture for Crop Wild Relatives conservation in Switzerland"

**Appendix S3** List of the available environmental variables available for SDMS. Each variable was assigned to a category: temperature (T), seasonality (S), extreme temperature (Tex), precipitation (P), aridity (A), topography (Topo), NDVI, orest height (Forest) and soil pH (pH). Each variable has a priority rank within its category for the preselection procedure. This procedure used a permutation test for which a minimum p-value (p-val th.) was required to keep the variable in the procedure.

| **Abbreviation** | **Description** | **Category** | **Priority** | **p-val th.** | **Source** |
| --- | --- | --- | --- | --- | --- |
| bio1_tmean | Annual mean temperature | T | 2 | 0.05 | ecospat |
| bio2_dr | Mean diurnal range | S | 4 | 0.05 | ecospat |
| bio3_iso | Yearly isothermality | S | 3 | 0.05 | ecospat |
| bio4_ts | Yearly temperature seasonality | S | 1 | 0.05 | ecospat |
| bio5_tmaxw | Max. temperature of the warmest month | Tex | 2 | 0.05 | ecospat |
| bio6_tminc | Min. temperature of the coldest month | Tex | 1 | 0.05 | ecospat |
| bio7_tar | Temperature annual range | S | 5 | 0.05 | ecospat |
| bio8_twetq | Mean temperature of the wettest quarter | Tex | 6 | 0.05 | ecospat |
| bio9_tdryq | Mean temperature of the dryest quarter | Tex | 5 | 0.05 | ecospat |
| bio10_twarmq | Mean temperature of the warmest quarter | T | 3 | 0.05 | ecospat |
| bio11_tcoldq | Mean temperature of the coldest quarter | T | 4 | 0.05 | ecospat |
| bio12_p | Annual precipitation | P | 1 | 0.05 | ecospat |
| bio13_pwet | Precipitation of the wettest month | P | 6 | 0.05 | ecospat |
| bio14_pdry | Precipitation of the driest month | P | 3 | 0.05 | ecospat |
| bio15_ps | Precipitation seasonality | S | 2 | 0.05 | ecospat |
| bio16_pwetq | Precipitation of the wettest quarter | P | 5 | 0.05 | ecospat |
| bio17_pdryq | Precipitation of the dryest quarter | P | 4 | 0.05 | ecospat |
| bio18_pwarmq | Precipitation of the warmest quarter | P | 2 | 0.05 | ecospat |
| bio19_pcoldq | Precipitation of the coldest quarter | P | 7 | 0.05 | ecospat |
| gdd3Y | Growing degree days | T | 1 | 0.05 | ecospat |
| arridity | Arridity (annual precipitation – potential evapotranspiration) | A | 1 | 0.1 | ecospat |
| sdiryy | Direct radiation | Topo | 3 | 0.1 | ecospat |
| slp25 | Slope | Topo | 2 | 0.1 | ecospat |
| topos | Convexity | Topo | 1 | 0.1 | ecospat |
| ndviMAX | Yearly maximum NDVI | NDVI | 3 | 0.1 | WSL |
| ndviMEAN | Yearly average NDVI | NDVI | 1 | 0.1 | WSL |
| ndviMIN | Yearly minimum NDVI | NDVI | 2 | 0.1 | WSL |
| ndviQ50 | Yearly median NDVI | NDVI | 5 | 0.1 | WSL |
| ndviQ80 | Yearly 80th percentile of NDVI | NDVI | 4 | 0.1 | WSL |
| ndviSD | Yearly NDVI standard deviation | NDVI | 6 | 0.1 | WSL |
| ForestQ25 | 25th percentile of the tree heights | Forest | 2 | 0.1 | WSL |
| ForestQ95 | 95th percentile of the tree heights | Forest | 1 | 0.1 | WSL |
| soilPH | Soil acidity | pH | 1 | 0.1 | WSL |

**Appendix S4. Species distribution modelling: detailed method**

We built species distribution models (SDMs) for all species with at least 10 spatially distinct occurrences (Appendix S5). We extracted the observations from the Info Flora database for all the priority CWR, with a minimal accuracy of 500 m. To reduce spatial autocorrelation, observations were disaggregated to keep a minimal distance of 300 m. SDMs relate species observation to environmental factors used as predictors. These environmental conditions were contrasted between species distributions and background data (or pseudo-absences) consisting of 10'000 points, randomly sampled throughout Switzerland. The initial set of predictors was composed of 33 variables available as georeferenced raster layers (Appendix S3). Bioclimatic variables are a fine scale resolution raster set produced by the ecospat laboratory at the University of Lausanne (Broennimann 2018). Soil, normalized difference vegetation index (NDVI) and forest canopy height were layers provided by the Swiss Federal Institute for Forest, Snow and Landscape Research (WSL). A detailed description of these layers can be found in the appendix 1 of Descombes et al. 2020.

A preliminary variable selection was processed for each taxon to reduce the number of predictors and avoid model overfitting. First, we applied a two-sample permutation test along each variable between occurrences points and the background points (Fay & Shaw 2010). Variables were retained only if there was a significant difference (Appendix S3 and S5). Then, we categorized these variables into 9 groups (temperature, precipitation, seasonality, extreme temperatures, aridity, topography, NDVI, soil pH and forest height; Appendix S3). We selected only one variable per group, based on the smallest p-value (i.e. the most discriminating variable). If p-values scores were ex aequo, we used a priority order based on our experience in SDMs to select the only one variable per group (Appendix S3). If a group showed no significant variable, no variable was retained for this group. After this initial step, the number of variables varied between two and 9 (i.e one per category) depending on the species.

The chosen modelling strategy depended on the number of observations per predictor. If this ratio was >= 10, three modeling algorithms were combined into an ensemble modeling approach (Thuiller et al 2004, Araùjo & New 2007). General Additive Models (GAM; Wood 2011), Maxent (ME; (Phillips et al. 2006; Elith et al. 2011; Merow et al. 2013) and Random Forest (RF; Liaw & Wiener 2002) were calibrated with the package *biomod2* (Thuiller et al. 2013, 2020; version 3.4.6). We used the default parametrization for GAM and RF but set the *beta multiplier* on 1.5 and did not use the *threshold feature* with ME. If the number of observations per predictor was <10, the modeling strategy consisted in an ensemble of small models (ESM; Breiner et al. 2015), using the same three algorithms. It consists in a combination of all the bivariate combinations of the retained predictors.

Models were evaluated with 4-fold cross-validation using 4 accuracy indices: the area under the curve (AUC; Pearce & Ferrier 2000), the true skill statistics (TSS_max_; Allouche et al. 2006) the sensitivity of the TSS_max_ and the continuous Boyce index (B; Hirzel et al. 2006). These 4 indices were scaled and combined into a consensus index, analogous to a correlation varying between -1 (total counter predictions) and 1 (perfect predictions), 0 meaning random predictions. The final predictions consist of the weighted mean of the predictions of each modelling technique. The consensus indices were used as weight and modelling techniques with a weight below 0.7 were not retained (Appendix S5). This threshold of 0.7 was also retained for the ESM: bivariate models with an average evaluation of 0.7 were not retained in the final assemblage of models.

Models were projected onto the set of environmental predictors, resulting in specie’s potential distributions. The initial continuous suitability maps scaled between 0 and 100 were reclassified into binarized predictions following 5 criteria: omission ratios of 5, 10 and 15, MaxKappa and AUC-optimized (Appendix S6 and S8; Q95, Q90, Q85, MaxKappa and RocOpt respectively). The omission ratios correspond to the percentile of training presences omitted from the potential distribution. An omission ratio of 10 means that 90% of the occurrences are included in the binarized distribution. The MaxKappa is the threshold maximizing the kappa metrics and the AUC-optimized threshold is the one minimizing the distance between the AUC-ROC plot and (0,1). Omission ratio thresholds were derived with a custom code (available on demand) wereahs MaxKappa and AUC optimized were obtained with the function *optimal.thresholds* from the *PresenceAbsence* R package version 1.1.9 (Freeman & Moisen 2008).

The R-code used to model the species distribution is available on demand to the authors.

***References***

Allouche O, Tsoar A, Kadmon R. 2006. Assessing the accuracy of species distribution models: Prevalence, kappa and the true skill statistic (TSS). Journal of Applied EcologyDOI: 10.1111/j.1365-2664.2006.01214.x.

Breiner FT, Guisan A, Bergamini A, Nobis MP. 2015. Overcoming limitations of modelling rare species by using ensembles of small models. Methods in Ecology and Evolution **6**.

Broennimann O. 2018. A high spatial and temporal resolution climate dataset for Switzerland. Page Technical Report. Available from www.unil.ch/ecospat/files/live/sites/ecospat/files/shared/PDF_site/chclim25.pdf (accessed August 31, 2022).

Descombes P, Walthert L, Baltensweiler A, Meuli RG, Karger DN, Ginzler C, Zurell D, Zimmermann NE. 2020. Spatial modelling of ecological indicator values improves predictions of plant distributions in complex landscapes. Ecography **43**.

Elith J, Phillips SJ, Hastie T, Dudík M, Chee YE, Yates CJ. 2011. A statistical explanation of MaxEnt for ecologists. Diversity and Distributions **17**.

Fay MP, Shaw PA. 2010. Exact and Asymptotic Weighted Logrank Tests for Interval Censored Data: The interval R Package. Journal of Statistical Software **36**:1–34. University of California at Los Angeles.

Freeman EA, Moisen G. 2008. PresenceAbsence: An R package for PresenceAbsence analysis. Journal of Statistical Software **23**.

Hirzel AH, le Lay G, Helfer V, Randin C, Guisan A. 2006. Evaluating the ability of habitat suitability models to predict species presences. Ecological Modelling **199**:142–152. Available from https://www.sciencedirect.com/science/article/pii/S0304380006002468 (accessed October 26, 2021).

Liaw A, Wiener M. 2002. Classification and Regression by randomForest. R News **2**.

Merow C, Smith MJ, Silander JA. 2013. A practical guide to MaxEnt for modeling species’ distributions: What it does, and why inputs and settings matter. Ecography **36**.

Pearce J, Ferrier S. 2000. Evaluating the predictive performance of habitat models developed using logistic regression. Ecological Modelling **133**.

Phillips SJ, Anderson RP, Schapire RE. 2006. Maximum entropy modeling of species geographic distributions. Ecological Modelling **190**:231–259. Available from https://www.sciencedirect.com/science/article/pii/S030438000500267X (accessed October 27, 2021).

Thuiller W, Georges D, Engler R. 2013. biomod2: Ensemble platform for species distribution modeling. R package version **2**.

Thuiller W, Georges D, Engler R, Breiner F. 2020. biomod2: Ensemble Platform for Species Distribution Modeling. Available from https://CRAN.R-project.org/package=biomod2.

Wood SN. 2011. Fast stable restricted maximum likelihood and marginal likelihood estimation of semiparametric generalized linear models. Journal of the Royal Statistical Society. Series B: Statistical MethodologyDOI: 10.1111/j.1467-9868.2010.00749.x.

**Appendix S8** Distribution of species richness based on the observations (Nsp_obs), the stack of binarized SDMs with an omission ratio of 5 (Nsp_OR5), the stack of binarized SDMs with an omission ratio of 10 (Nsp_OR10), the observations after quantile normalization (Nsp_obs_std), the stack of binarized SDMs with an omission ratio of 10 after quantile normalization (Nsp_OR10_std), the stack of binarized SDMs with an omission ratio of 15 (Nsp_OR15), the stack of binarized SDMs with the best kappa (Nsp_kappa), the stack of binarized SDMs based on the AUC optimisation (Nsp_roc), the stack of binarized SDMs with an omission ratio of 15 after quantile normalization (Nsp_OR15_std), the stack of binarized SDMs with the best kappa after quantile normalization(Nsp_kappa_std), the stack of binarized SDMs based on the AUC optimisation after quantile normalization(Nsp_roc_std)

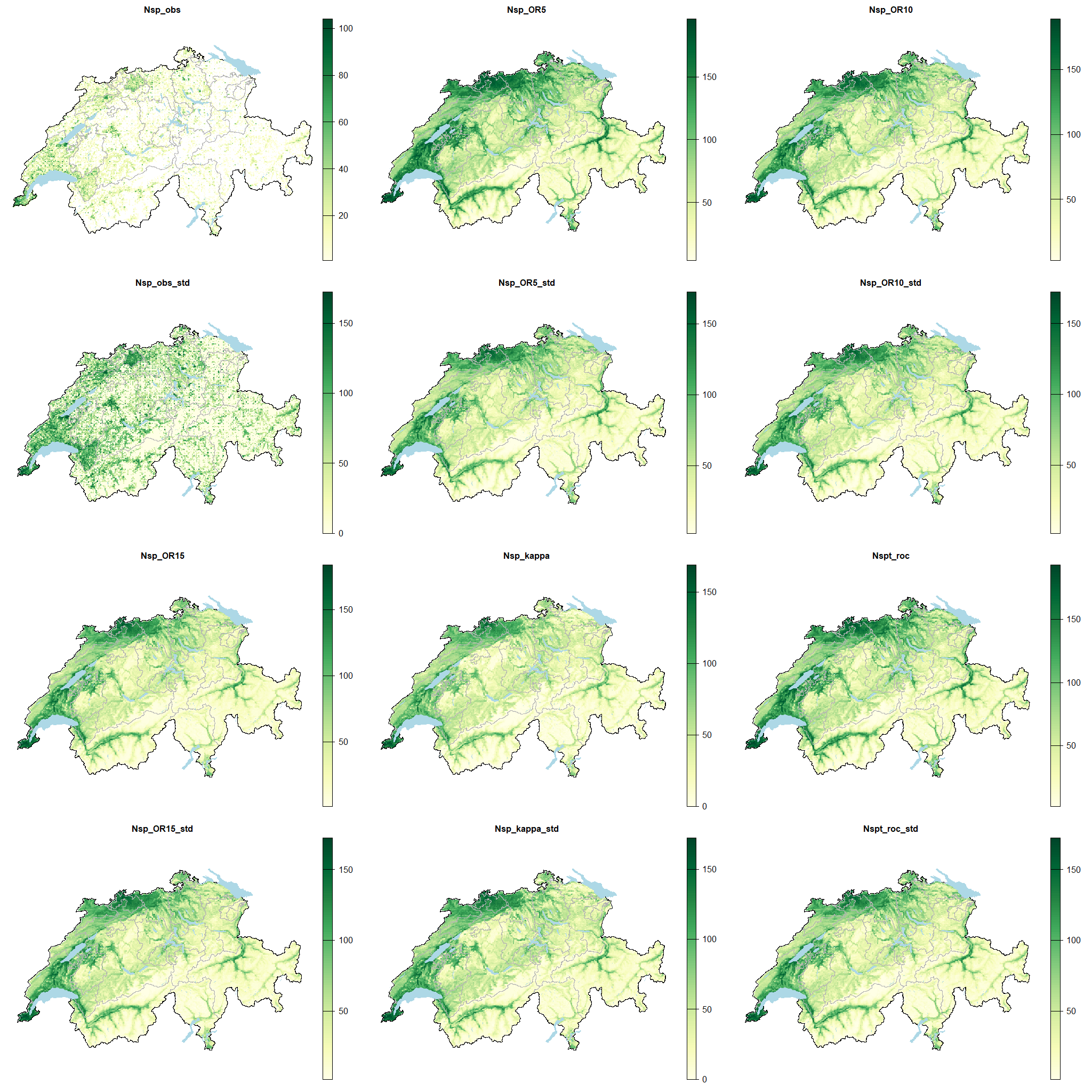
